# Double knockout CRISPR screen in cancer resistance to T cell cytotoxicity

**DOI:** 10.1101/2022.03.01.482556

**Authors:** Jonathan J. Park, Adan Codina, Lupeng Ye, Stanley Lam, Jianjian Guo, Paul Clark, Xiaoyu Zhou, Lei Peng, Sidi Chen

## Abstract

Immunotherapy has transformed cancer treatments; however, a large fraction of patients encounter resistance. Such resistance is mediated by complex factors, often involving interactions between multiple genes. Thus, it is crucially important to identify genetic interactions between genes that are significantly mutated in cancer patients and those involved in immune responses, ideally the ones that currently have chemical compounds for direct targeting. To systematically interrogate such genetic interactions that mediate cancer cells’ response to T cell killing, we designed an asymmetric CRISPR/Cas9 dual perturbation library targeting the matched combinations between significantly mutated tumor suppressors and immune resistance genes. We performed a combinatorial double knockout screen on 1,159 gene pairs and identified those where joint loss-of-function renders altered cellular response to T cell cytotoxicity. With individual double knockout constructs we validated these genetic interactions including *Jak1-Trp53, Jak1-Kmt2d*, and *Ifngr1-Kmt2d*. Interactions between significantly mutated tumor suppressors and potentially druggable immune resistance genes may offer insights on potential new concepts of how to target clinically relevant cancer mutations with currently available agents. This study also provides a technology platform and an asymmetric CRISPR double knockout library for interrogating genetic interactions between cancer mutations and immune resistance pathways under various settings.

## Background

Despite impressive durable responses elicited by cancer immunotherapy, the majority of patients do not see long-term benefit with treatment ^1,2^. This has motivated many ongoing studies on identification of both tumor cell-intrinsic and extrinsic factors that affect the immunogenicity of tumors ^3–7^, however the molecular mechanisms that determine therapeutic resistance remain poorly understood. In particular, most genetic correlates of immune responsiveness to date have been made with single genetic events rather than with cooperative genetic interactions ^8,9^, with notable exceptions including the immunosuppressive phenotype being driven by cooperativity between Myc overexpression and Ras mutation ^10^, as well as *STK11/LKB1* mutation in KRAS-driven lung adenocarcinoma ^11^. Such synergistic oncogenic alterations may play an important role in response to immunotherapy but are difficult to identify due to lack of experimental tools for studying functional consequences of gene interactions at scale.

One recent strategy to enable scalable systematic mapping of genetic-interaction networks is to adapt the CRISPR-Cas9 screening technology for massively parallel knockout of single genes and gene pairs ^12,13^. Simultaneous gene perturbation can be used to study the coordinated behaviors of genes and, in particular, whether the phenotypic effects of mutations are different when acting in concert rather than individually. While CRISPR-Cas9-based screens have been used to identify single gene events that contribute to resistance to immune cell killing ^14,15^, there has been limited studies using combinatorial CRISPR screens for identifying novel cooperative gene interactions contributing to immune cytotoxicity resistance in an unbiased, high-throughput manner. Identifying such interactions may generate deeper insights into the underlying biological networks that drive tumor-cell intrinsic mechanisms of immunotherapy resistance as well as potentially identify genetic vulnerabilities for combinatorial immune-based therapies.

## Methods

### Cell lines

Single cell clones were derived from the murine melanoma cell line (B16F10) transduced with PGK-mCherry-OVA lentivirus to reduce cellular heterogeneity and ensure homogenous expression of antigen. Single cells were sorted and multiple clones were derived. A clonal cell line (clone #3) was used for subsequent experiments. A combination of antibiotic selection and flow cytometry was used to ensure purity. These cells were subsequently transduced with Cas9 lentivirus to create B16F10;OVA;Cas9 clone #3 cells (referred to as BC3 cells). All screens were conducted in this background. BC3 cells were transduced with Firefly Luciferase (FLuc) lentivirus for bioluminescence assays, and all experiments were conducted in the BC3-FLuc background. All cell lines were grown under standard conditions using D10 (DMEM supplemented with 10% FBS, 1% Penicillin-Streptomycin) in an incubator maintained at 37 °C with 5% CO_2_.

### Design of CADRE library

The Combinatorial Antineoplastic Drug Resistance Experiment (CADRE) library follows an asymmetric design, combining the most significantly mutated tumor suppressors in human cancers with immunotherapy resistance associated genes derived from the antigen processing and presentation, IFN-gamma, MAPK, PI3K/AKT/MTOR, and WNT/beta-catenin signaling pathways. To generate the immunotherapy resistance gene set, the union of the following gene lists were used “regulation_of_MAP_kinase_activity” (GO: 0043405), “antigen_processing_and_presentation” (GO: 0019882), “interferon-gamma-mediated_signaling_pathway” (GO: 0060333), “T_cell_costimulation” (GO: 0031295), “HALLMARK_PI3K_AKT_MTOR_SIGNALING” (MSigDB: M5923), “HALLMARK_WNT_BETA_CATENIN_SIGNALING” (MSigDB: M5895), “HALLMARK_INTERFERON_GAMMA_RESPONSE” (MSigDB: M5913), with the following gene lists removed “T_cell_receptor_complex” (GO: 0042101), “B_cell_receptor_signaling_pathway” (GO: 0050853), “Toll-like_receptor_1-Toll-like_receptor_2_protein_complex” (GO: 0035354), “Toll-like_receptor_2-Toll-like_receptor_6_protein_complex” (GO: 0035355). This gene list was then intersected with the a list of drug targets obtained by cross-reference of the UniProtKB/Swiss-Prot manually reviewed, complete human proteome (proteome identifier up000005640) with the DrugBank database of targets for FDA-approved small molecule drugs, FDA-approved biotech drugs, nutraceuticals, and experimental drugs. The tumor suppressor gene list was curated from Kandoth et al., 2013 ^16^. sgRNAs were designed using a custom pipeline integrating sgRNA quality scoring by CRISPOR ^17^. In total, the library consists of 1,159 DKO gene pairs represented by 8,321 DKO sgRNA-sgRNAs; 632 SKO genes represented by 1,684 sgRNA-NTCs; and 84 DNTCs. The library was synthesized as an oligo pool (CustomArray).

### Lentiviral production and transduction

Prior to transduction media was removed from 80-90 percent confluent HEK293FT cells and replaced with OptiMEM serum free media to enhance transduction efficiency. Envelope packaging plasmid pMD2.G, packaging plasmid psPAX2, and dual-sgRNA lentiviral plasmids were combined in a ratio of 1:1.5:2 and suspended in OptiMem serum free media. Polyethyleneimine (PEI) was added to plasmid DNA pool (8 μL 1mg/mL PEI per 1μg DNA) and mixed gently before being incubated at room temperature for 15 min. After incubation, plasmid solutions were added dropwise to HEK293FT cells. 6 hours post transduction OptiMEM media was replaced with D10 media. Supernatant was collected from HEK293FT cells after 48 h post transduction and spun down at 1000 x *g*, 4 °C, for 5 min to remove cellar debris. Viral supernatant was then aliquoted before being frozen at -80 °C prior to experimentation. To determine viral titer to be infected cells were counted and seeded in plates at appropriate densities, before being infected with various dilutions of viral supernatant. 24 h post infection titrated puromycin was added to infected cells (10 μg/mL) and cultured for 72 h. Cell survival was then assayed to determine the functional viral titer of the supernatant.

### Double knockout cellular library production

Each screen was performed with four infection replicates, low multiplicity of infection (MOI), and high screening coverage. Briefly, BC3 cells were seeded at a density of 5e6 cells per plate in 15cm plates were transduced with 1e6 functional viral particles per plate for a calculated MOI of 0.2, and incubated for 24 h prior to replacing media with fresh media containing 4 μg/mL puromycin for selection. A total of 2.5e7 cells (5 plates of 5e6) were seeded and approximately a total of 5e6 cells were infected, conferring ∼500x library coverage.

### Naïve OT-I CD8^+^ T cell isolation and culture

Mouse CD8^+^ T cell isolation and culture methods were based on our previous works ^18,19^. Briefly, mesenteric lymph nodes (mLNs) and spleens were dissected from OT-I mice, then placed into ice-cold PBS supplemented with 2 % FBS. Lymphocytes were released by grinding organs through a 100 μm filter, then re-suspended with 2 % FBS. Red blood cells (RBCs) were lysed with ACK lysis buffer (Lonza). RBC-lysed lymphocyte solution was filtered through 40 μm filters to remove cell debris. Naïve CD8a^+^ T cell purification was performed using Naïve CD8a^+^ T cell Isolation Kits (Miltenyi Biotec) according to the manufacturer’s protocols. Naïve CD8a^+^ T cells were cultured in RPMI-1640 (Gibco) media supplemented with 10 % FBS, 2 mM L-Glutamine, 200 U / mL penicillin–streptomycin (Gibco), and 49 μM β-mercaptoethanol (Sigma), hereafter referred to as cRPMI media. Naïve CD8a^+^ T cells were activated with anti-CD3ε and anti-CD28 antibodies (BioLegend), For *in vitro* experiments, cRPMI media was supplemented with 2 ng / mL IL-2, 1 μg / mL anti-CD28, and 12 ng / mL Il-12p70 cytokines or antibodies. All cytokines and antibodies were purchased from BioLegend. For antigen stimulation co-culture experiments OT-1 cells were collected as described above, but activated by culturing harvested T-cells with OVA expressing cancer cells in a 1:1 ratio. Antigen stimulated OT-1 cells were activated for 3-4 days prior to co-culture experimentation.

### Asymmetric CRISPR double knockout screen

Library transduced cells resuspended in cRPMI were seeded at a density of 2.5e4 into three 96 well plates to achieve a coverage of ∼720X for the R1 screen per E:T ratio. OT-I CD8^+^ T-cells were added to each well at either E:T 2 or 5 and cultured for 48 h before the introduction of 1ug/ml puromycin to remove selective pressure of T-cells. Cells were then allowed to rest for 72 h before being collected for gDNA extraction. The screen was repeated in entirety using the same methodology to have independent experimental biological replicates, with the sole exception of using 6 plates per E:T ratio to increase library coverage.

### Genomic DNA extraction

To isolate gDNA from CADRE screen cells, cells were washed three times with PBS to remove cellular debris from dead cells before being collected and pooled by condition. Pooled cells were spun down at 400 x *g* for 5 min and reconstituted in 200 μL of PBS before being extracted with Qiagen blood mini gDNA extraction kit according to manufacturer’s protocol. To isolate gDNA for non-screen samples ∼2.5e4 cells were taken per sample, and washed with PBS. PBS was aspirated and resuspended in 100 μL Lucigen QuickExtract buffer and incubated for 30 minutes at 65 °C before being heat inactivated at 95 °C for 5 min.

### CADRE library readout

Two rounds of PCR were used for sgRNA library readout. PCR #1 used genomic DNA (∼2 μg per reaction and 3 reactions per sample) for sufficient coverage of screen, and PCR #2 used 1μL of PCR#1 product with barcoded primers. Samples were amplified with different barcoded primers and pooled for deep sequencing.

The following cycle parameters were used for PCR #1: 98 °C for 1 min, 25 cycles of (98 °C for 1 s, 62 °C for 5 s, 72 °C for 15 s). The following primers were used as well:

Forward: 5’-aatggactatcatatgcttaccgtaacttgaaagtatttcg-3’

Reverse: 5’-ctttagtttgtatgtctgttgctattatgtctactattctttccc-3’

The following cycle parameters were used for PCR #2: 98 °C for 30 s, 28 cycles of (98 °C for 1 s, 62 °C for 5 s, 72 °C for 15 s), and 72 °C 2 min for the final extension. See Table S4 for barcoded primers. PCR reactions were performed using Phusion Flash High Fidelity Master Mix (ThermoFisher). Gel purification of pooled products from a 2% E-gel EX (Life Technologies) were performed using the QiaQuick Gel Extraction kit (Qiagen).

### CADRE library mapping

Raw paired FASTQ files were filtered and demultiplexed using Cutadapt ^20^. To demultiplex the barcodes in the forward PCR primers used during readout, the following settings were used cutadapt -g file:fbc.fasta –no-trim, where the fbc.fasta contained the forward barcodes. To pare down the forward read to the first 20 base pair sgRNA spacer sequence, and the reverse read to the second spacer sequence, the following settings were used cutadapt -g GTGGAAAGGACGAAACACCG -G CTCTAAAAC -l 20 -e 0.2 -m 19 –discard-untrimmed. The reverse complement of the second spacer sequence from the reverse read was obtained using fastx_reverse_complement from FASTX-Toolkit, and then combined with the first spacer sequence from the forward read to create 40bp fused spacer-spacer sequences. These 40bp fused sequences were then mapped to the fused sgRNA sequences from the CADRE library (Supplementary Table 2) for dual sgRNA quantification using BWA-ALN ^21^. A BWA index of the sgRNA library was generated using the bwa index command, and SAMTools was used for post-processing ^22^.

### Minimum count threshold

We determined a minimum count threshold used for downstream analysis (**Figure S1D**). First, we measured frequencies of the SKO sgRNAs in the library, calculated the expected frequencies of the double sgRNAs, and compared them to observed DKO double sgRNAs frequencies. The ratio of observed to expected frequencies fell markedly below a read count of 15. The sgRNAs with counts below this threshold (read count of 15) were then masked from further analysis.

### Identification of gene interaction sgRNA pairs

We used two methods to identify potential gene interactions. Sample counts were read normalized to 1e6. In order to see if the phenotypic effect of DKO dual gene pair perturbation was different from the constitutive SKO perturbations, we performed the two-sided Wilcoxon rank sum test on DKO sgRNAs abundances compared to gene A and gene B SKO sgRNA abundances for gene pairs. In order to determine enrichment-based gene interactions, we calculated the observed and expected DKO enrichment:

E_observed_ = (D_s_/D_c_) * (N_c_/N_s_)

E_expected_ = (A_s_/A_c_) * (N_c_/N_s_) + (B_s_/B_c_) * (N_c_/N_s_)

D_s_ = median abundance for DKO pair observed in post-selection screen samples

D_c_ = median abundance for DKO pair observed in pre-T cell treatment cell control samples

N_c_ = median abundance for DNTCs observed in pre-T cell treatment cell control samples

N_s_ = median abundance for DNTCs observed in post-selection screen samples

A_s_ = median abundance for SKO gene A observed in post-selection screen samples

A_c_ = median abundance for SKO gene A observed in pre-selection cell control samples

B_s_ = median abundance for SKO gene B observed in post-selection screen samples

B_c_ = median abundance for SKO gene B observed in pre-selection cell control samples

To determine the outlier gene interactions, linear regression was performed on E_expected_ vs E_observed_ using the lm function in R, and the outlier.test function from the “car” package was used to determine Bonferroni p-values for the most extreme observations.

### Luciferase assay for cell survival

Luciferase readout was done both 24 and 48 h after co-culture in 96-well white polystyrene plates. 150 μg/mL D-luciferin (PerkinElmer) was added using a multichannel pipette to cells, and covered from light and incubated for 10 min. After 10 min, luciferase intensity was measured using a plate reader (PerkinElmer)

### TCGA transcriptome analyses

For global comparative TCGA gene expression profile analyses, Gene Expression Profiling Interactive Analysis (GEPIA) ^23^ and GEPIA2 ^24^ were used. Log normalized transcripts per million (TPM) values were visualized for normal RNA-seq samples from GTEx and tumor RNA-seq samples from TCGA across 33 different cohorts or selected cohorts using individual gene queries or gene signature queries, with significance thresholds set at |log2FC| cutoff of 0.5 and q-value cutoff at 0.01. Correlation analyses used Spearman’s coefficient. Cell type proportion analysis of GTEx normal and TCGA tumor samples were performed using GEPIA2021 and CIBERSORT for deconvolution.

For additional *KMT2D* correlation analysis, skin cutaneous melanoma (SKCM) TCGA RNA-seq samples were obtained from the Broad GDAC and normalized to TPM. Spearman correlations were calculated using the cor function in R. Genes that were significantly positively or negatively correlated with *KMT2D* were determined using a cutoff to select approximately the top or bottom 10% of the sorted values, and the identified genes were used for Database for Annotation, Visualization and Integrated Discovery (DAVID)^25,26^ functional gene annotation analysis.

### Survival analyses

Survival analyses based on the expression status of genes were performed using GEPIA2 and parameters Group Cutoff: Median, Cutoff-High(%): 50; Cutoff-Low(%): 50. Survival maps were also created, using a significance level of 0.05. Survival analyses for how query genes affect the influence of cytotoxic T lymphocyte (CTL) levels on patient outcomes were performed using the Tumor Immune Dysfunction and Exclusion (TIDE) algorithm ^27^.

### Gene mutation profiles in patient cohorts

*KMT2D, JAK1, JAK2, IFNGR1*, and *TP53* allele frequencies was queried using cBioPortal ^28,29^. All melanoma studies were selected for visualization and analyses, leading to a combined study of 2834 samples from 2781 patients in 15 studies. Alteration frequencies were obtained using a minimum number of total cases threshold of 10, and mutual exclusivity and co-occurrence analyses were obtained for all pairwise combinations. Statistics were determined by cBioPortal.

## Results

It is crucially important to identify genetic interactions between two key partners – (1) tumor suppressors that are significantly mutated in cancer and thereby have direct clinical relevance, and (2) immunotherapy resistance associated genes that are potentially targetable with small molecule inhibitors or chemical compounds. Interactions between tumor suppressors and druggable immune resistance targets can offer insights on new concepts of how to target clinically prevalent cancer mutations with currently available agents. To systematically interrogate such genetic interactions that mediate immune resistance, we employed an asymmetric, combinatorial CRISPR Cas9 screening approach. We designed a Combinatorial Antineoplastic Drug Resistance Experiment (CADRE) screening strategy with an asymmetric library design, combining the pan-cancer most significantly mutated tumor suppressors (“TS genes”) in human cancers on one side of the combination; with putative drug targetable immunotherapy resistance-associated genes (“IR genes”) on the other, which include the antigen processing and presentation, IFN-gamma, MAPK, PI3K/AKT/MTOR, and WNT/beta-catenin signaling pathways (**Figure 1A-B**). The CADRE library targets the combinations between 61 immune resistance genes and 19 tumor suppressors across cancer types, with a total of 1,159 genetic combinations (gene pairs) (**Figure 1B**). For each gene pair, 3 guides were chosen in most cases, for a total of 9 sgRNA pairs of double knockout (DKO) constructs per gene pairs for most gene pairs (except certain cases where optimal sgRNAs were not available). In addition, corresponding single knockout (SKO) constructs were represented in the library, as well as double non-targeting controls (DNTCs) for a total of 10,089 constructs (**Figure 1B**). All guides were selected on the basis of sgRNA quality scoring by CRISPOR ^17^ including cutting efficiency, out-of-frame patterns, and specificity (**Figure S1B**).

**Figure 1.**
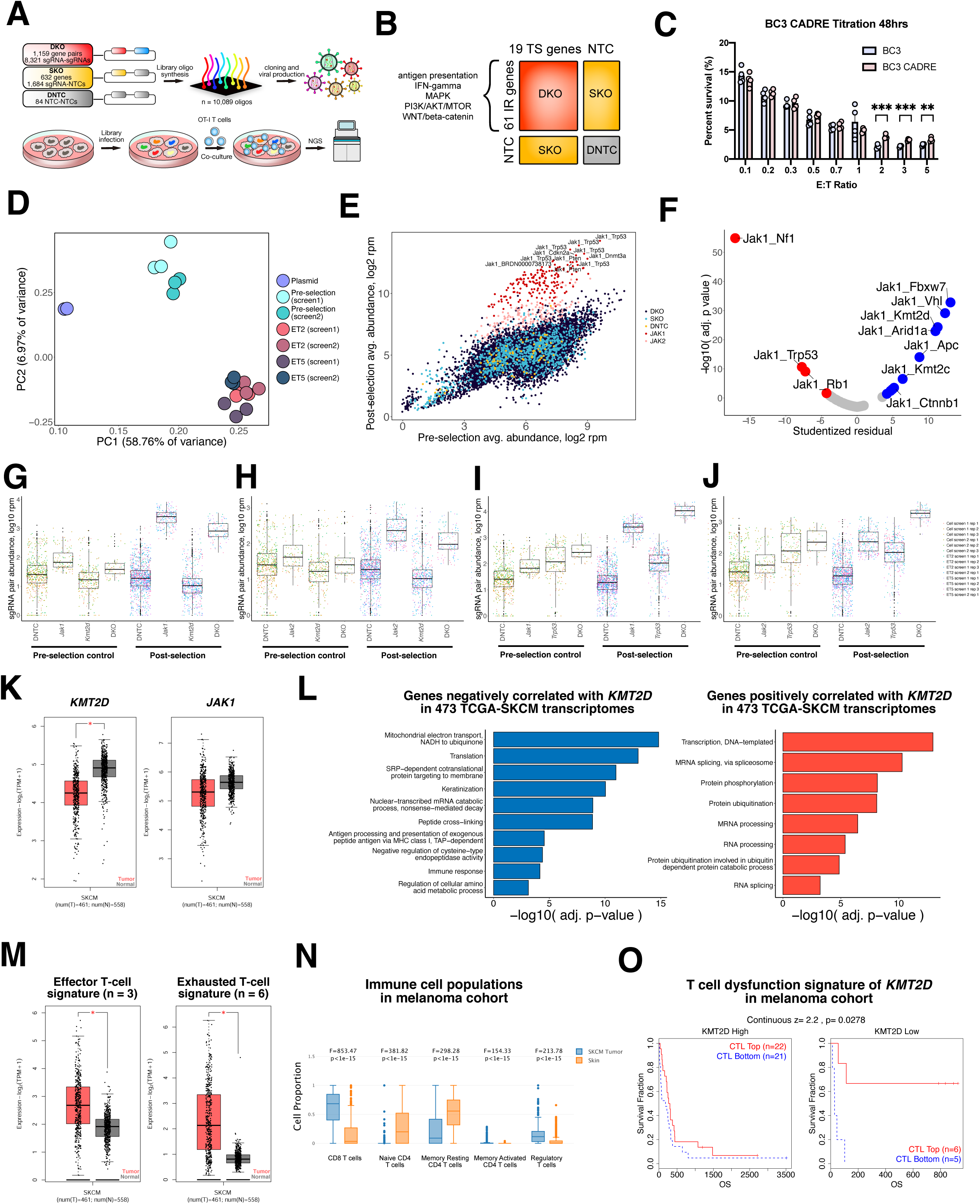
Asymmetric double knockout CRISPR screen of gene pairs that affect cancer cell response to T cell killing. (A) Schematic overview of CADRE screen. (B) Schematic of CADRE library design, 61 genes with immunotherapy resistance were crossed in combinatorial fashion with 19 significantly mutated tumor suppressors to create a DKO pool. SKOs and NTCs serve for comparison and as controls. (C) Titration of BC3 cells, and BC3 CADRE cells co-cultured with E:T ratios ranging from 0.1 to 5. High E:T ratios demonstrated significant phenotypic differences and were therefor selected for screening (q values of 1.76e-3 and 1.19e-2 by multiple T test, 1% FDR for E:T ratios 2 and 5 respectively). *, adjusted p-value < 0.05. **, adjusted p-value < 0.01. ***, adjusted p-value < 0.001. (D) Principle component analysis (PCA) of the sgRNA pair read count distributions across screens, E:T ratios, technical replicates, and pre-T cell treatment controls. (E) Scatterplots comparing guide representation of the CADRE library in post co-culture samples averaged across all replicates, E:T ratios, and screens compared to pre-selection infected cell controls. *Jak1* and *Jak2* associated sgRNA pairs (either DKO or SKO) are marked in red. (F) Scatterplot comparing Bonferroni adjusted p value determined by outlier test compared to Studentized residuals from linear regression analysis in (C). (G-J) Tukey box plots (IQR boxes with 1.5 × IQR whiskers) overlaid on dot plots of sgRNA pair abundances for each DKO, SKO, and DNTCs for pre-T cell treatment controls (also labelled as “cell”) and post-selection co-culture samples with reads pooled from samples across screens, E:T ratios, and technical replicates. Count distributions are shown for gene pairs (G) *Jak1_Kmt2d*, (H) *Jak2_Kmt2d*, (I) *Jak1_Trp53*, and (J) *Jak2_Trp53*. (K) Boxplots of *KMT2D* and *JAK1* expression in RNA-seq samples from the TCGA SKCM melanoma cohort and paired normal samples (461 tumor samples, 558 normal samples). *, q-value < 0.01 and |log2FC| > 0.5. (L) Bar plots of top enriched pathways identified by DAVID biological processes analyses of genes negatively (left) and positively (right) correlated with *KMT2D* expression in 473 melanoma RNA-seq samples from TCGA. (M) Boxplots of effector T-cell gene signatures (*CX3CR1, FGFBP2, FCGR3A*) and exhausted T-cell gene signatures (*HAVCR2, TIGIT, LAG3, PDCD1, CXCL13, LAYN*) in RNA-seq samples from the TCGA SKCM melanoma cohort and paired normal samples (461 tumor samples, 558 normal samples). *, q-value < 0.01 and |log2FC| > 0.5. (N) Boxplots of normalized cell type proportions from CIBERSORT deconvolution analyses of TCGA-SKCM and GTEx normal skin RNA-seq samples for T cells. Statistics shown on plots. (O) Kaplan-Meier curves showing the survival of melanoma patients from the GSE22153 cohort based on the expression status of *KMT2D* and linked to estimated cytotoxic T lymphocyte (CTL) levels. Analyses performed with the Tumor Immune Dysfunction and Exclusion (TIDE) algorithm. Statistics shown on plot.

The CADRE library was synthesized via oligonucleotide array and cloned into lentiviral vectors (**Figure S1A**), and the representations of DKOs, SKOs and DNTCs in the library were verified by next-generation sequencing (NGS) (**Figure S1C-D**). While every single guide in the library (10,089 / 10,089) was found to have some read representation in the plasmid samples, we found that below a read count of 15 the ratio of observed to expected DKO sgRNA frequencies fell markedly (**Figure S1D**). Guides with counts below this threshold (15 reads) were masked from further analysis yielding 10,031 sgRNAs. CADRE lentiviral library was generated and transduced into B16F10 melanoma cells stably expressing antigenic ovalbumin (OVA) protein, Cas9 protein, and reporter firefly luciferase (FLuc). Murine B16F10 melanoma cells were chosen as a model due to their resistance to immune checkpoint blockade against receptors such as programmed cell death protein 1 (PD-1) and cytotoxic T-lymphocyte-associated protein 4 (CTLA-4) ^30,31^. OVA protein transduction was performed due to availability of transgenic mice with OT-I T cells that recognize OVA peptide of amino acids 257–264 (SIINFEKL) expressed in the context of H-2Kb ^32,33^, which allows for antigen-specific T cell killing assays. Lentiviral infection occurred at a low multiplicity of infection (MOI) (MOI < 0.2) to bias towards single lentiviral sgRNA construct integration. Additionally, lentiviral library infection occurred at a coverage of approximately 500X to ensure high representation of the sgRNA pairs in the CADRE library. We NGS verified that the transduced pre-selection cell pool retained the vast majority of the DKO, SKO and DNTC constructs in the CADRE library, with 10,029 / 10,031 sgRNAs detected in the pre-selection samples (**Figure S1E**).

We then assayed the innate resistance of these cells by co-culturing library and non-library infected B16F10;OVA;Cas9 clone #3 cells (hereafter referred to as **BC3** cells) with OT-I CD8+ T-cells which homogeneously recognize SIINFEKL epitope presented by the cell line. A titration series of effector to target ratios (E:T ratios) were assayed ranging from 0.1 T-cells per cancer cell to 5 T-cells per cancer cell. Survival was measured by bioluminescence assay at 24 and 48 hours to determine cancer cell survival. Across E:T ratios, library and non-library transduced cell survival was comparable, with the exception of the high E:T ratio conditions (E:T ratios > 1) where the mutant pool demonstrated a significant increase in resistance compared to baseline (**Figure 1C**). Given the increased resistance of library transduced cells at higher E:T ratios, E:T ratios of two and five were chosen to conduct the screen. We co-cultured BC3-CADRE cells with OT-I CD8+ T-cells in 96 well plates, and 72 hours post selection low dose puromycin (1ug/ml) was introduced to culture conditions to remove T-cell selective pressure and allow surviving cells to rest. After 72 hours of resting, plates were washed with PBS, and cells were collected for genomic extraction. This screening process was repeated as a whole to generate independent biological experimental replicates for more robust analysis. The sgRNA cassettes were PCR amplified from each condition’s gDNA across both screens for both post-selection samples and pre-selection cell pool controls, barcoded, and deep-sequenced by NGS, which revealed high level coverage of the CADRE library in all samples sequenced (**Figure S2A**). Clustering analysis of the library representation showed distinct clusters between plasmid, cell population before co-culture, and cell populations post co-culture (**Figure S2B**), suggesting a high-quality screen and NGS readout. Principal components analysis using the mapped read counts revealed consistency between the cell pool conditions and the E:T ratios two and five across both screens (**Figure 1D**). Based on the cumulative distribution functions (CDFs) of the representative samples, there are strong shifts between pre-selection and post-selection co-cultures (**Figure S2C**), indicative of strong selection seen at sgRNA library levels.

Using DNTC sgRNA pairs to represent an empirical null distribution without selection, at a false-discovery rate (FDR) of 1.19% we identified 222 enriched sgRNA pairs of which 194 (87.4%) are associated with *Janus kinase 1 (Jak1)* or *Janus kinase 2 (Jak2)*, including DKO and SKO constructs. Bulk analysis revealed that *Jak*-associated sgRNAs dominated the enrichment in the screen post-selection (**Figure 1E; Figure S3A-D**). In the context of tumor immune resistance, *Jak1* LOF mutations have consistently demonstrated a functional role in resistance to checkpoint blockade immune therapy ^34^. The dominance of *Jak1/2* in the enriched resistant population proved the robustness of the T cell resistance phenotype measuring the effect of the screen.

We then compared the representation of the dual-sgRNAs of each DKO gene pair with the representation of the SKOs of their constitutive genes, and found that *Jak1/2*-associated gene pairs were the most statistically significantly different from their constitutive SKOs (**Figure 1E-F**), suggestive of potential gene interactions. In order to determine whether such gene interactions may be additive or subtractive, we calculated phenotype scores for observed DKO enrichment compared to expected DKO enrichment (SKO gene A enrichment + gene B enrichment), normalized to DNTCs and pre-T cell treatment cell control representation. While the immune cell killing phenotype can be reliably predicted as sums of individual gene perturbations, using the Bonferroni Outlier Test for extreme values from the linear regression and comparing the values against the Studentized residuals, we observe that gene pairs *Jak1_Trp53, Jak1_Nf1, and Jak1_Rb1* have higher observed enrichment for double knockout than expected (Bonferroni-adjusted p-value < 0.001) suggesting potential additive gene interaction (**Figure 1F**), while gene pairs *Jak1_Apc, Jak1_Vhl, Jak1_Kmt2c, Jak1_Kmt2d, Jak1_Arid1a, Jak1_Fbxw7, Jak1_Ctnnb1* (Bonferroni-adjusted p-value < 0.001) have lower observed enrichment for double knockout than expected, suggesting potential subtractive gene interaction (**Figure 1F**). Boxplots of normalized read counts for *Jak1_Kmt2d, Jak2_Kmt2d, Jak1_Trp53* and *Jak2_Trp53* (**Figures 1G-J**) suggest subtractive and additive phenotypic interactions to *Jak1/2* perturbation for *Kmt2d* and *Trp53*, respectively. *Jak1*, the dominant gene found in our analysis, is a non-receptor tyrosine kinase associated with interferon gamma (IFN-gamma) signaling. While this role of *Jak1* has been linked to immunotherapy resistance ^34^, which support the rigor of the screen, it was still surprising to see the significant interaction between *Jak1* and *Kmt2d*. KMT2D (Lysine Methyltransferase 2D) is an epigenetic modifier that regulates transcription of multiple pathways including beta-globin and estrogen receptor genes. KMT2D has also been shown to be a tumor suppressor gene in multiple human cancers ^16,35–38^.

In order to further study the significance of the genes of interest identified in this study and how they may potentially interact, we have performed a series of comparative transcriptomics-based analyses on tumor and normal samples from the Cancer Genome Atlas (TCGA) and Genotype-Tissue Expression (GTEx), both across many TCGA cancer types and for the skin cutaneous melanoma (SKCM) cohort specifically. First we looked at the global gene expression profiles of *KMT2D, JAK1, TP53*, and *IFNGR1* across all tumor samples and paired normal tissues (**Figure S4A-D**) and more specifically for *KMT2D* and *JAK1* in the SKCM cohort (**Figure 1K-M**), and identified tumor type specific expression patterns. We found that KMT2D expression levels were significantly decreased in melanoma samples compared to normal tissue, likely due to its tumor suppressor functionality.

Next we looked at the expression levels of gene signatures in the SKCM cohort and found that the effector and exhaustion T-cell signatures were upregulated in the tumor samples (**Figure 1M**). Moreover, cell proportion deconvolution analyses with CIBERSORT revealed increased estimated proportions of CD8 T cells, memory activated CD4 T cells, and Tregs in the tumor samples, with a decrease of naïve and memory resting CD4 T cells (**Figure 1N**). These results suggest a tumor microenvironment where there is both increased infiltration of effector CD8 T cells albeit with concomitant increased exhaustion and immunosuppression. The increased infiltration of CD8 T cells along with the association of high tumor mutational burden and neoantigens with better responses to immunotherapy ^6,39^ highlight the importance of MHC class I based antigen processing and presentation in tumor-immune interactions for certain tumor types such as melanoma and non-small cell lung cancer.

To further evaluate the significance of *KMT2D* and *JAK1* in clinical cohorts, we performed correlation analyses to determine which genes and signatures were associated with *KMT2D* expression levels. We calculated the Spearman correlations for all genes compared to *KMT2D* expression in the TCGA SKCM melanoma cohort and identified positively and negatively correlated genes. Genes negatively correlated with *KMT2D* were further analyzed using DAVID gene ontology functional annotation and found to be enriched for mitochondrial electron transport, translation, peptide cross-linking, antigen processing and presentation via MHC-I, and immune response (**Figure 1L**). Genes positively correlated with *KMT2D* were found to be enriched in transcription, mRNA splicing, protein phosphorylation, mRNA processing, and ubiquitination. These results suggest that *KMT2D* has an important role in MHC-I based antigen processing and presentation, which is consistent with our experimental data and model whereby *KMT2D* loss may potentially increase antigen presentation. We also find that specific class I MHC genes are also negatively correlated with *KMT2D* expression: Spearman coefficients for HLA-A is - 0.093165455; HLA-B, -0.062233376; HLA-C, -0.047136103, which provides further evidence for this model. We also performed correlation analyses between *KMT2D* and *JAK1* expression and genes associated with the ontology term “interferon-gamma mediated signaling pathway” (GO ID 0060333) to see if there is any relationship between *KMT2D* and IFN-gamma signaling. We found positive and significant correlations for both *JAK1* and IFN-gamma signaling gene signatures across both the SKCM cohort and across 33 different cancer types from TCGA (**Figure S5A**).

In addition to examining the transcriptional profiles of the genes of interest from this study, we looked at the gene mutation profiles and frequencies across several melanoma studies (combined study of 2834 samples from 2781 patients in 15 studies, using cBioPortal) to ensure that the identified genes have clinical significance at the genomic level. We found that *KMT2D* and *JAK1* are both frequently mutated in melanoma patients (up to ∼30% and ∼15% alteration frequencies respectively, **Figure S5B**). Mutual exclusivity and co-occurrence analyses for all pairwise combinations of *KMT2D, JAK1, JAK2, IFNGR1*, and *TP53* suggest that all mutation combinations except *JAK2-IFNGR1* co-occur at a significant rate (**Figure S5C**), suggesting that these may have some clinical significance and can be leveraged for patient stratification.

We also performed survival analyses on the TCGA-SKCM patient cohort and the pan-cancer TCGA cohort to see the effect of expression status of *KMT2D* and *JAK1* on patient survival (**Figures S4E-F, S5D-E**). We found that for patients from the SKCM cohort, those with high KMT2D expression had significantly worse disease free survival (hazard ratio 1.3, p = 0.026) whereas high JAK1 expression had no effect on survival. For pan-cancer analysis, both high KMT2D and JAK1 expression led to a decreased hazard ratio for disease free survival. Survival maps (**Figures S4G-H**) revealed cancer type specific effects of *KMT2D, JAK1, JAK2, IFNGR1*, or *TP53* expression levels on patient survival. These results suggest that for melanoma patients, decreased *KMT2D* expression may lead to improved survival, perhaps in part due to increased antigen presentation, but the survival effects do not necessarily generalize to all cancer types. Given the high variability regarding immune infiltration, tumor mutational burden, and the genomic profiles for different cancer types, cancer type specific patterns are to be expected.

We also performed Tumor Immune Dysfunction and Exclusion (TIDE) analysis ^27^ and examined the survival of melanoma patients from the GSE22153 cohort ^40^ based on the expression status of *KMT2D* and linked to estimated cytotoxic T lymphocyte (CTL) levels. The *KMT2D*-low patient group demonstrated increased CTL-associated overall survival benefit, whereas high levels of *KMT2D* abolished the overall survival benefit of CTL-high patients (**Figure 1O**). These results suggest that *KMT2D* deficiency is linked to the effect of infiltrating CTL on melanoma patient survival.

## Discussion

Genetic interactions occur when perturbation of genes in the same or related pathways result in phenotypes that differ from the sum of the effects of the individual mutations ^41–43^. Studying such interactions provides insight into on the coordinated behavior of genes and potentially identify new therapeutic avenues; for example, drugs targeting poly(ADP-ribose) polymerase (PARP) enzymes have shown some efficacy for *BRCA1/2* mutant tumors, representing a synthetic lethal strategy that incorporates the patient’s genetic background for targeted precision medicine ^44^. However, tools for identifying such interactions at scale, particularly for complex phenotypes such as immunotherapy resistance, have thus far been limited.

CRISPR–Cas9 systems offer scalable and precise gene editing with minimal off-target effects, allowing for adaptations into high-throughput screening strategies for dissection of functional gene interaction networks. Recent examples of such strategies have been in models of cell fitness or sensitivity to small molecule inhibitors ^12,13^. However, we were interested in developing a combinatorial screening platform for a more complex tumor-immune model. Our group has previously performed a screen of significantly mutated genes (SMGs) in human cancers using CRISPR-mediated genetically engineered mouse models (CRISPR-GEMM) in immune checkpoint blockade (ICB) settings ^45^. The strength of this system is that GEMMs can more precisely mimic features of human cancers as the tumors develop *in vivo* in a preserved immune microenvironment. The limitations of this model include the time scale of the experiments (on the order of months) and that although the screening vector has an additional sgRNA cassette targeting *Trp53*, the library design is still single gene KO and limited in scale (49 SMGs). Likewise, other tumor-immune CRISPR screens performed to date ^14,15^ have also been based on single gene perturbation. The goal and novelty of our platform was for truly combinatorial (e.g. double gene KO) screening in an antigen-specific *in vitro* co-culture model. Although such models are not as sophisticated for recapitulating the tumor microenvironment as *GEMMs*, their advantages include: (1) the time scale of the assays are much faster, on the order of days, (2) larger scale for library design, in this case over a thousand gene pairs, (3) isolation of selective pressure, in this case specifically for T cell mediated killing, and (4) simplicity of the co-culture system for ease of adaption to other tumor cell types and immune cell types.

We have also performed an extensive set of analyses on clinical cohorts which in aggregate demonstrate that for melanoma patients, *KMT2D* is significantly negatively correlated with antigen presentation via MHC-I, is correlated with IFN-gamma signaling signatures, has significantly high co-occurrence with *JAK1* at the mutational level, and is associated with decreased survival in TCGA patients with low *KMT2D* levels improving the survival of CTL-high patients based on TIDE analysis. These human cancer patient based genomic data and analyses are consistent with our murine model based experimental results which suggest that *KMT2D* LOF can attenuate immune resistance for mutations in the interferon gamma signaling pathway potentially through rescue of antigen presentation. Further study will be necessary to definitively identify how these processes occur on a molecular basis. Overall, we provide here evidence which sheds light on the potential interactions between *JAK1* and *KMT2D*, and have demonstrated the potential of double knockout screening as a tool for discovery in cancer immunology.

The double knockout screen system here is able to identify pairs whose simultaneous LOF would render tumor cells very sensitive to T cell killing, potentially identifying drugs which may potentiate the effect of immunotherapy in conjunction with the presence of biomarkers. We unexpected found *Jak1/Jak2* pairs to dominate the pool of the top hits, which limited the number of other gene pairs without these two genes. *KMT2D*, for example, currently has no specific inhibitors available. The double knockout screen and the resulting gene pairs identified in the study have clinical relevance. First, the unbiased double knockout screen demonstrated the observation that JAK1/2 is a dominating signal cancer resistance to T cell killing in a pool double mutant setting of competition. Second, the screen results identified and validated gene pairs mediating cancer resistance, in a quantitative manner. Third, these gene pairs provided potential double-biomarkers for patient stratification to improve the probability of immunotherapy efficacy. Moreover, we believe that the dual perturbation CRISPR screening strategy can serve as a starting point to better understand immunotherapy resistance at a systems level, as we have shown with comparative analyses in human patient cohorts. Finally, the double knockout approach provided here can be used in other contexts and with other library designs to identify new therapeutic avenues with more success.

## Conclusions

Here, we developed a high-throughput CRISPR-based dual perturbation genetic screening strategy with asymmetric library design and antigen-specific tumor-immune interaction models to identify genetic interactions underlying cellular response to T cell killing between cancer mutations and potentially druggable immune resistance pathways. The most striking phenotype observed in our screens was the effect that *Jak1 / Jak2* mutations had on tumor cell resistance to cytotoxic effector T cell killing. Loss-of-function mutations in these genes have been associated with anti– programmed death 1 (PD-1) therapy resistance in patient populations ^34^. Genetic interaction analyses revealed that *Kmt2d* seemed to have a buffering effect and *Trp53* a synergistic effect on the *Jak1* phenotype. Moreover, *Jak1* and *Kmt2d* mutations significantly co-occur in a multitude of human cancers, including melanoma (cBioPortal mutual exclusivity analysis for pan cancer studies q = <0.01), suggesting our asymmetric screen approach can identify clinically relevant genetic interactions. These findings are consistent with our previous study, where we found that *KMT2D* deficiency sensitizes tumors to immune checkpoint blockade in GEMMs ^45^, and expands on our knowledge of *Kmt2d* deficiency as a tumor cell intrinsic vulnerability predisposing factor in context of combinatorial co-mutations. Together, we demonstrate how dual loss-of-function CRISPR screens with asymmetric library designs can be extended to complex phenotypes such as resistance to immune cell killing. Such approaches may help adapt precision medicine approaches to extend the striking efficacy of immunotherapy to a broader population.

## List of abbreviations

BC3 cells: B16F10;OVA;Cas9 clone #3 cells
FLuc: firefly luciferase
CADRE: Combinatorial Antineoplastic Drug Resistance Experiment
SMG: significantly mutated gene
LOF: loss of function
GEMM: genetically engineered mouse model
CTL: cytotoxic T lymphocyte
ICB: immune checkpoint blockade
TIDE: Tumor Immune Dysfunction and Exclusion
DAVID: Database for Annotation, Visualization and Integrated Discovery
TCGA: The Cancer Genome Atlas
SKCM: skin cutaneous melanoma
GTEx: Genotype-Tissue Expression
OVA: ovalbumin
DKO: double knockout
SKO: single knockout
NTC: non-targeting control
DNTC: double non-targeting control

## Declarations

### Ethics approval and consent to participate

This study has received institutional regulatory approval. All recombinant DNA and biosafey work was performed under the guidelines of Yale Environment, Health and Safety (EHS) Committee with an approved protocol (Chen-rDNA-15-45; Chen-rDNA-18-45). All animal work was performed under the guidelines of Yale Institutional Animal Care & Use Committee (IACUC) with an approved protocol (Chen-2018-20068).

### Consent for publication

Not applicable.

### Availability of data and materials

All data generated or analyzed during this study are included in this article and its supplementary information files. Specifically, source data and statistics for non-high-throughput experiments are provided in Supplemental Tables. Processed data for high-throughput sequencing experiments are provided as processed quantifications in Supplemental Datasets. Raw sequencing data are being deposited to NIH Sequence Read Archive (SRA) or Gene Expression Omnibus (GEO) with pending accession number(s). Original cell lines are available at commercial sources listed in supplementary information files. Genetically modified cell lines are available via Chen lab. Most data, reagents, methods, computational codes and materials that support the findings of this research are available from the corresponding author upon reasonable request. Some material used in the reported research may require requests to collaborators and agreements with other entities. Requests are reviewed by Yale University to verify whether the request is subject to any intellectual property or confidentiality obligations. Any material that can be shared will be released via a Material Transfer Agreement.

## Authors’ contributions

JJP conceived and designed the study, developed computational pipelines and analyzed the data. JJP, AC and LY performed experiments. SL, JG, PC, XZ, and LP assisted with various experiments. SC conceived the study, provided conceptual advice, secured funding, and supervised the work. JJP, AC, LY, and SC wrote the manuscript. All authors read and approved the final manuscript.

## Acknowledgments

We thank all members in Chen laboratory, as well as various colleagues in Department of Genetics, Systems Biology Institute, Cancer Systems Biology Center, MCGD Program, Immunobiology Program, BBS Program, Cancer Center, Stem Cell Center, Liver Center, RNA Center and Center for Biomedical Data Sciences at Yale for assistance and/or discussion. We thank the Center for Genome Analysis, Center for Molecular Discovery, High Performance Computing Center, West Campus Analytical Chemistry Core, and Keck Biotechnology Resource Laboratory at Yale, for technical support.

## Sample size determination

Sample size was determined according to the lab’s prior work or similar studies in the literature.

## Randomization and blinding statements

Regular *in vitro* experiments were not randomized or blinded. High-throughput experiments and analyses were blinded by barcoded metadata.

## Standard statistical analysis

Standard statistical analyses were performed using regular statistical methods. GraphPad Prism, Excel and R were used for analyses. Different levels of statistical significance were accessed based on specific p values and type I error cutoffs (0.05, 0.01, 0.001, 0.0001). Details of statistical tests were provided in supplemental information.

## Code availability

Codes used for data analysis or generation of the figures related to this study are available from the corresponding author upon reasonable request.

## Figure legends

**Figure S1.**
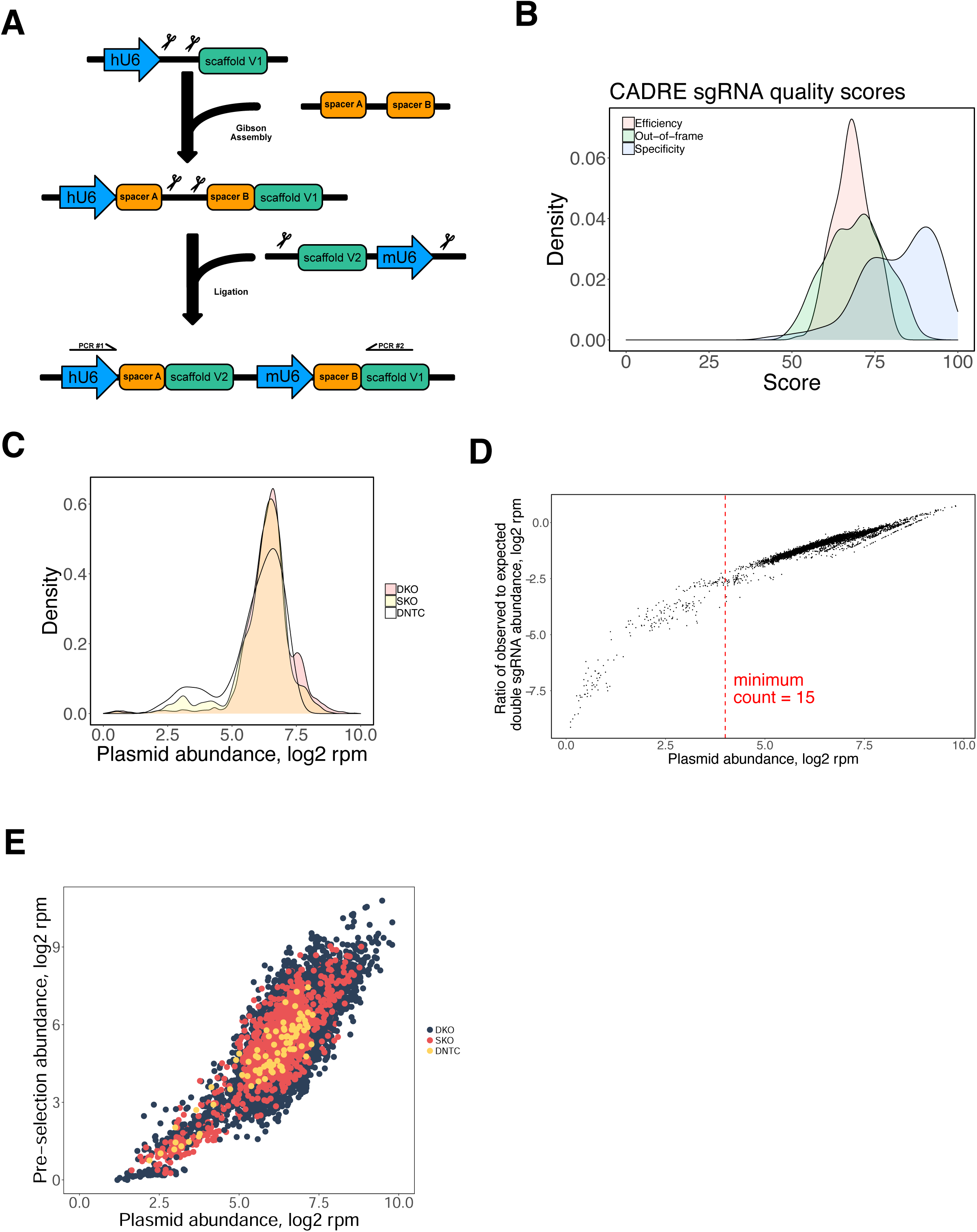
Construction and sequencing of the CADRE CRISPR knockout library in plasmid and transduced cell pools. (A) Schematic overview of double perturbation CRISPR construct designs. Long oligonucleotides containing paired guide sequences were synthesized and cloned into a lentiviral vector. Subsequently, a gene fragment containing a secondary sgRNA scaffold and mU6 promoter were cloned in between the paired guide sequences to reconstitute two fully functional sgRNA expression systems. (B) Density plot representing the distribution of scores for cutting efficiency, out-of-frame patterns, and specificity for sgRNAs comprising the CADRE library. (C) Density plot for the sgRNA pair representation in the CADRE plasmid library. (D) Minimally required sgRNA pair read count estimation from plasmid library. Observed frequency of double sgRNA was compared to expected frequency calculated from single sgRNAs reads. This analysis revealed that below ∼15 read counts, this ratio falls below expected. Guides with less than 15 read counts were masked from further analysis. (E) Scatterplot comparing guide representation of the CADRE library in the pre-T cell treatment infected cell controls averaged across all replicates and screens to the plasmid control.

**Figure S2.**
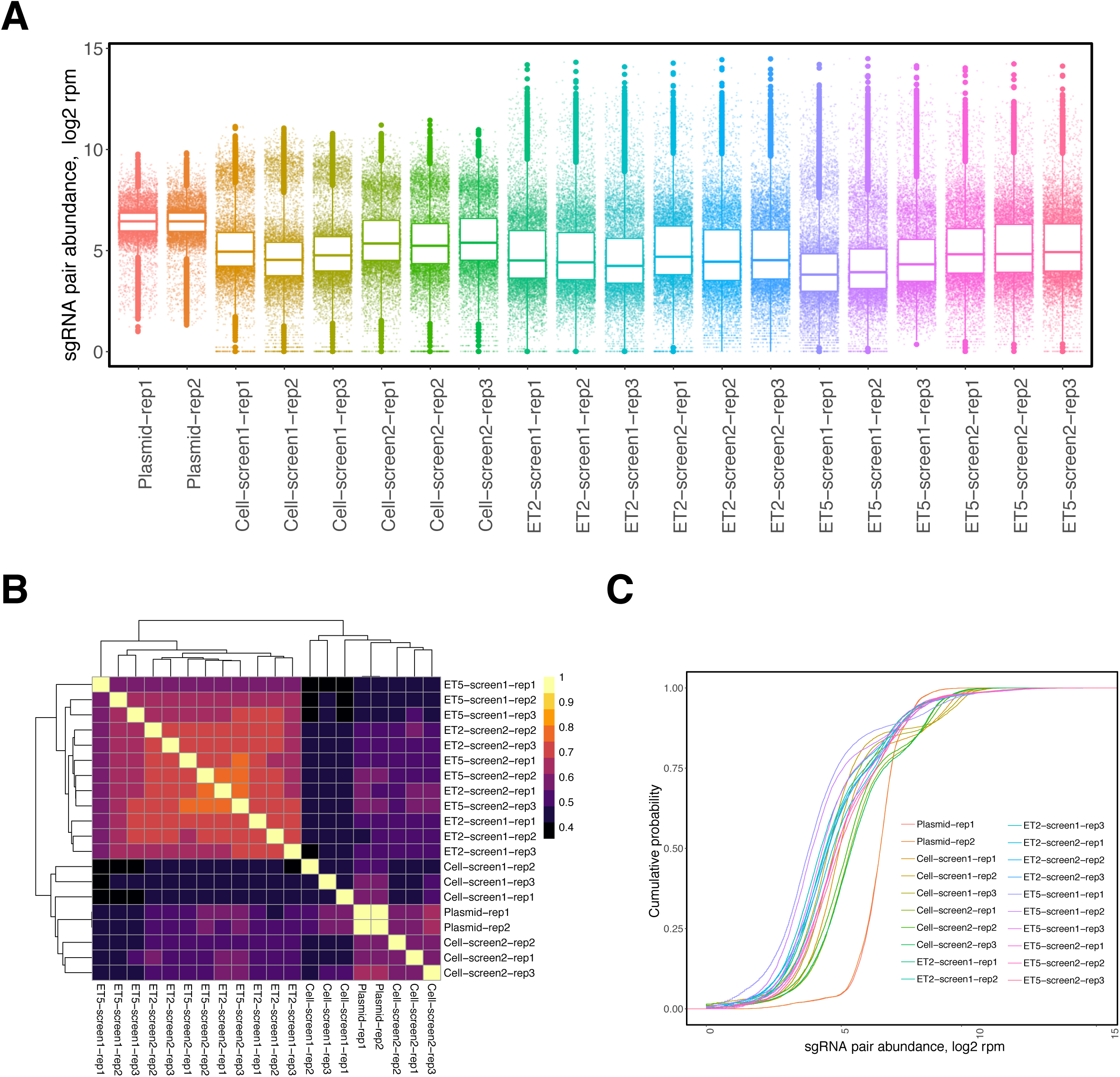
CADRE asymmetric double knockout CRISPR screen revealed dynamic cellular population shifts before and after co-culture selection. (A) Tukey box plots (IQR boxes with 1.5 × IQR whiskers) overlaid on dot plots of sgRNA pair abundance in samples across screens, E:T ratios, technical replicates, and both pre-T cell treatment (labelled as “cell”) and plasmid controls. (B) Heatmap showing Pearson correlation of log normalized sgRNA pair abundances across samples. (C) Empirical cumulative distribution function (CDF) plot of sgRNA pair abundances across samples.

**Figure S3.**
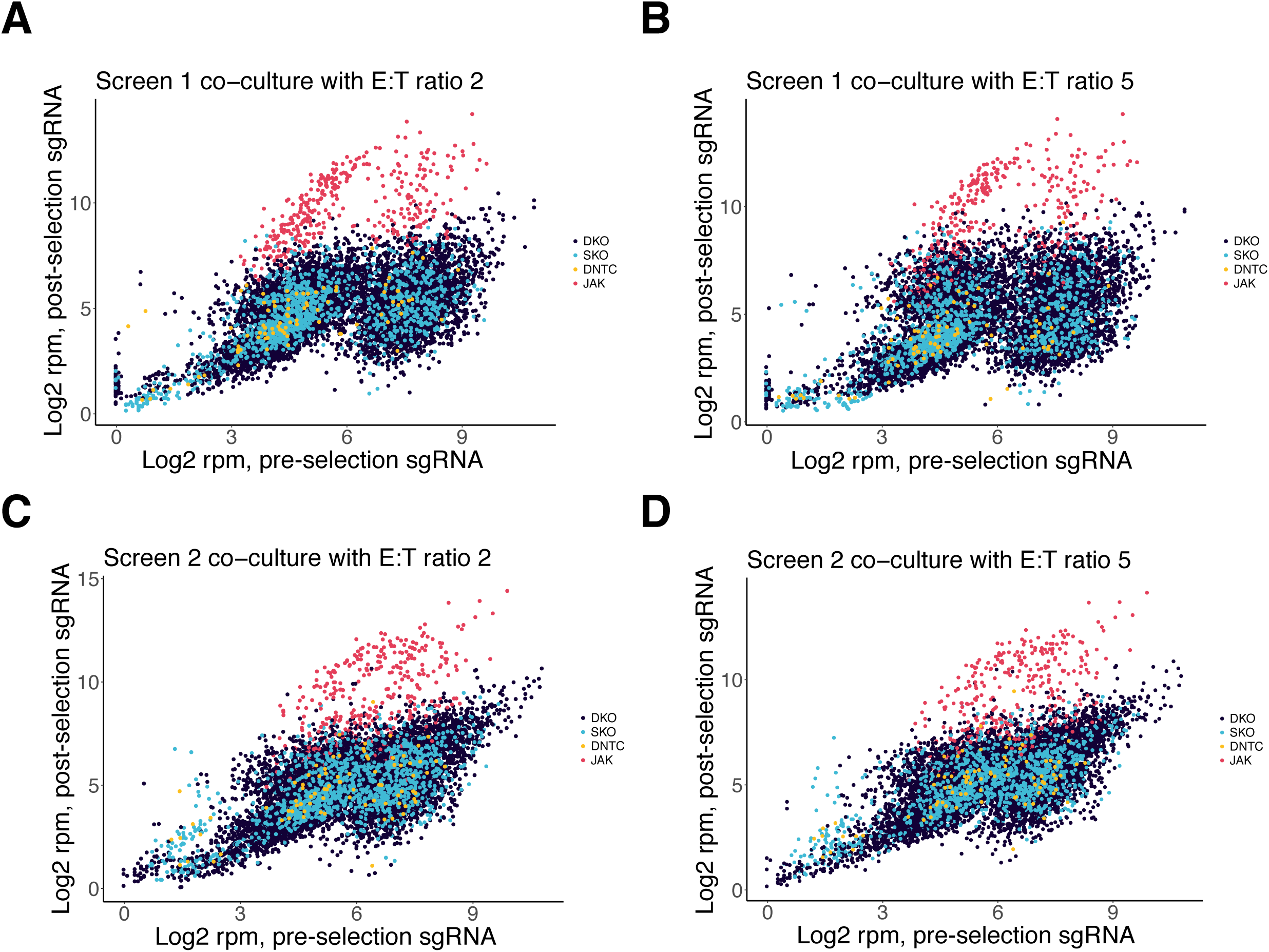
Additional analysis of CADRE double knockout CRISPR screen on T cell killing. (A–D) Scatterplots comparing guide representation of the CADRE library in post co-culture samples compared to pre-T cell treatment infected cell controls for (A) screen 1 and E:T ratio 2, (B) screen 1 and E:T ratio 5, (C) screen 2 and E:T ratio 2, (D) screen 2 and E:T ratio 5. *Jak1* and *Jak2* associated sgRNA pairs (either DKO or SKO) are marked in red (JAK), with other non-JAK DKO sgRNA pairs marked in dark blue, other SKO sgRNA pairs marked in light blue, and DNTCs marked in yellow.

**Figure S4.**
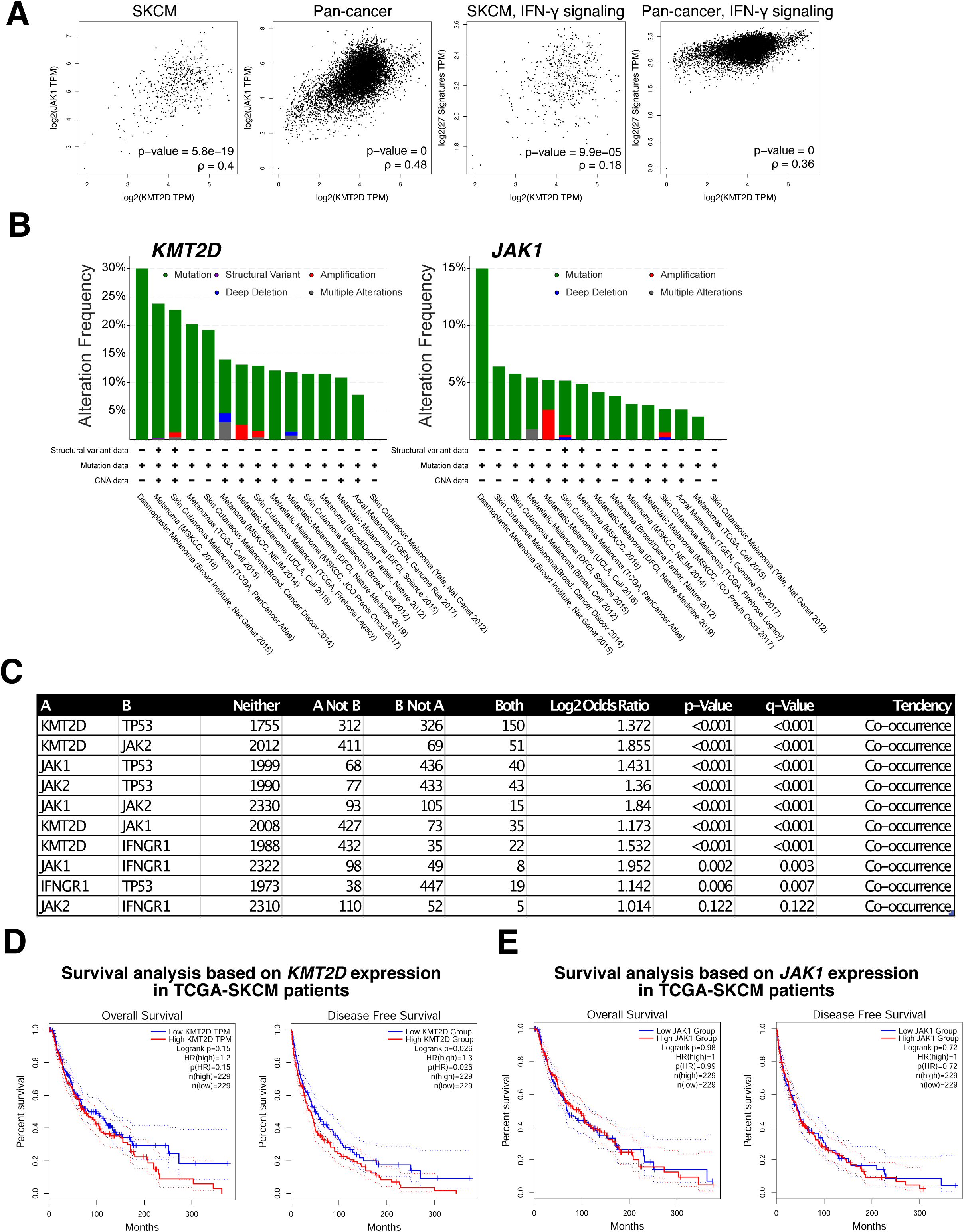
Additional analyses of TCGA cancer and matched normal RNA-seq samples. (A–D) Dot plots showing the gene expression profiles of (A) *KMT2D*, (B) *JAK1*, (C) *TP53*, and (D) *IFNGR1* across all tumor samples and paired normal tissues. Green dots represent normal samples, red dots represent tumor samples. Cohorts with q-value < 0.01 and |log2FC| > 0.5 are labelled in green if expression levels are greater for the normal samples, and in red if levels are greater for the tumor samples. (E) Kaplan-Meier curves showing the survival of patients from 33 different TCGA cohorts based on the expression status of *KMT2D*. Statistics shown on plot. (F) Kaplan-Meier curves showing the survival of patients from 33 different TCGA cohorts based on the expression status of *JAK1*. Statistics shown on plot. (G) Survival map showing the overall survival contributions of *KMT2D, JAK1, JAK2, IFNGR1*, and *TP53* across multiple TCGA cohorts. (H) Survival map showing disease free survival contributions of *KMT2D, JAK1, JAK2, IFNGR1*, and *TP53* across multiple TCGA cohorts.

**Figure S5.**
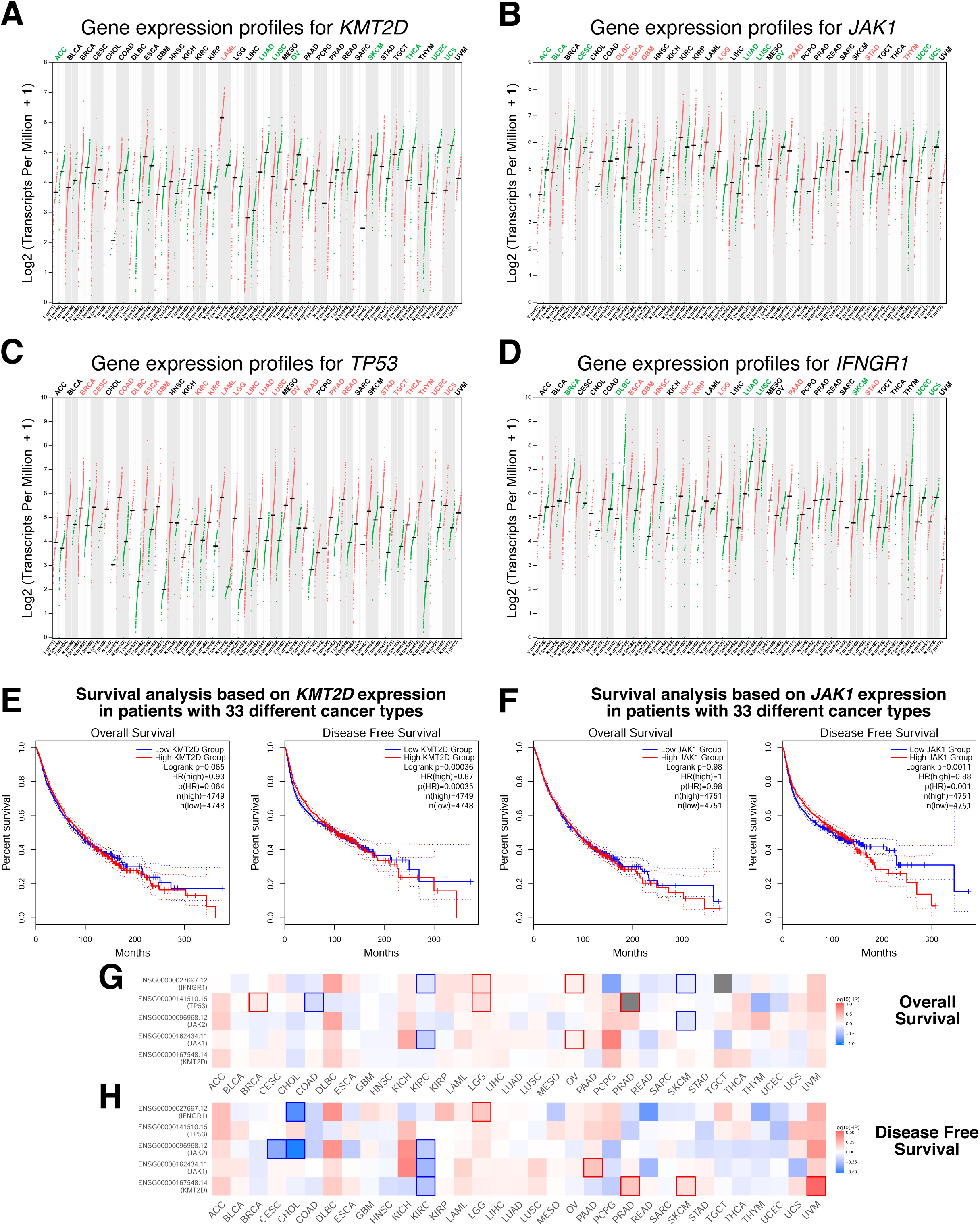
Analyses of TCGA cancer and matched normal RNA-seq samples and mutational profiles. (A) Scatterplots comparing *KMT2D* expression with (far left) *JAK1* expression in TCGA SKCM dataset; (left) *JAK1* expression in pan-cancer TCGA dataset of 33 types; (right) interferon-gamma-mediated signaling pathway gene signature (gene ontology 0060333) in TCGA SKCM dataset; (far right) interferon-gamma-mediated signaling pathway gene signature in pan-cancer TCGA dataset of 33 types. (B) Alteration frequencies for *KMT2D* and *JAK1* in 2834 samples from 15 melanoma studies. (C) Co-mutation analyses for *KMT2D, JAK1, JAK2, IFNGR1*, and *TP53* across melanoma studies. Statistics shown in table. (D) Kaplan-Meier curves showing the survival of TCGA-SKCM patients based on the expression status of *KMT2D*. Statistics shown on plot. (E) Kaplan-Meier curves showing the survival of TCGA-SKCM patients based on the expression status of *JAK1*. Statistics shown on plot.

## List of Supplemental Tables (provided in a compound excel file)

S1. CADRE dual sgRNA library by gene pair.

S2. Concatenated sgRNA sequences for mapping CADRE.

S3. SgRNA quality scoring by CRISPOR.

S4. Second PCR NGC barcoded readout primers.

S5. CADRE abundance counts for plasmid, cell control, and screens.

S6. Wilcoxon rank sum test significance for DKO dual sgRNA construct abundance compared to SKO gene A abundance or SKO gene B abundance.

S7. Observed DKO dual sgRNA construct enrichment compared to expected enrichment determined by SKO gene A enrichment + SKO gene B enrichment.

S8. Studentized residual vs significance from gene pair outlier test on linear fit of observed vs expected DKO enrichment.

S9. Spearman correlation values for comparisons to *KMT2D* with gene expression values from TCGA SKCM melanoma samples.

S10. DAVID analysis for genes negatively correlated with *KMT2D* in TCGA SKCM samples. S11. DAVID analysis for genes positively correlated with *KMT2D* in TCGA SKCM samples.

